# A Comparison of Skeletal Muscle Diffusion Tensor Imaging Tractography Seeding Methods

**DOI:** 10.1101/2024.08.29.610343

**Authors:** Bruce M. Damon, Roberto Pineda Guzman, Carly A. Lockard, Xingyu Zhou

**Author notes:** Address for Correspondence: Bruce M. Damon, PhD, Stephens Family Clinical Research Institute Carle Foundation Hospital, 611 W. Park Ave. Urbana, IL 61801.

## Abstract

The internal arrangement of a muscle’s fibers with respect to its mechanical line of action (muscle architecture) is a major determinant of muscle function. Muscle architecture can be quantified using diffusion tensor magnetic resonance imaging-based tractography, which propagates streamlines from a set of seed points by integrating vectors that represent the direction of greatest water diffusion (and by inference, the local fiber orientation). Previous work has demonstrated that tractography outcomes are sensitive to the method for defining seed points, but this sensitivity has not been fully examined. To do so, we developed a realistic simulated muscle architecture and implemented four novel methods for tract seeding: seeding along the muscle-aponeurosis boundary with an updated procedure for rounding seed points prior to lookup in the muscle boundary mask and diffusion tensor matrix (APO-3); voxel-based seeding throughout the muscle volume at a user-specified spatial frequency (VXL-1); voxel-based seeding throughout the muscle volume at a variable spatial frequency (VXL-2), and seeding near external and internal muscle boundaries (VXL-3). We then implemented these methods in an example human dataset. The updated aponeurosis seeding procedures allow more accurate and robust tract propagation from seed points. The voxel-based seeding methods had quantification outcomes that closely matched the updated aponeurosis seeding method. Further, the voxel-based methods can accelerate the overall workflow and may be beneficial in high throughput analysis of multi-muscle datasets. Continued evaluation of these methods in a wider range of muscle architectures is warranted.

## 1. Introduction

Skeletal muscle architecture, defined here as the number and geometric properties of a muscle’s fibers with respect to its mechanical line of action, impacts several functional aspects of skeletal muscle contraction. For example, muscle fiber length and orientation impact muscle force production, length excursion, and shortening or lengthening velocity^1–3^, while muscle fiber curvature may influence the magnitudes and spatial patterns of intramuscular fluid pressure^2,4,5^ and strain^6^ development. Brightness-mode ultrasound^7^ and diffusion-tensor MRI (DT-MRI) tractography^8^ have been used to study muscle architecture. Because MRI can also spatially map several of the functional properties of muscle contraction, there is special interest in developing DT-MRI tractography as a quantitatively accurate method for studying muscle architecture and its relationship to function.

It is well established that water diffusion in skeletal muscle is anisotropic, with the direction of greatest diffusion aligned with the muscle fiber’s long axis^9^. In the diffusion tensor model, the direction of greatest diffusion is represented by the tensor’s first eigenvector. Deterministic tractography algorithms integrate these vectors from an array of seed points, resulting in streamlines that represent the local muscle architecture on the spatial scale of several fascicles. How accurately these streamlines represent the fascicles’ geometry depends on several data acquisition and analysis conditions. Recent works have explored the impact and optimal selections for slice thickness, in-plane voxel size, signal-to-noise ratio (SNR), integration method, step-size, termination criteria, and tract post-processing methods, arriving at a set of proposed best practices^10–14^.

The present work continues this examination, with the specific focus being the definition of seed points. Two general approaches to seed point definition are described in the literature. First, tracts may be seeded at or near to the boundary of the muscle with its internal aponeurosis^15^. The advantage of this approach is that it facilitates calculations of mechanically relevant structural properties, such as the angle formed by the fibers and the aponeurosis (often reported as the pennation angle and symbolized as θ or α; but here symbolized as γ to avoid confusion with other angles of interest). As reported here, the aponeurosis-based seeding approach also allows estimation of the angles formed by 1) the muscle’s fibers and its mechanical line of action and 2) the aponeurosis and the muscle’s mechanical line of action (α and β, respectively, as defined in Ref.^16^). To define the aponeurosis geometry, manual definition^15^ or automated selection based on signal thresholds^17^ or diffusion properties^18^ can be used. A disadvantage to aponeurosis-based seeding is that partial volume artifacts, digitization errors, or rounding conventions may require the seeding mesh to be dilated away from the true muscle-aponeurosis boundary. This may result in fiber tract lengths that underestimate the true fascicle length. As discussed in Ref.^19^, other seeding approaches include placing seed points in the voxels of specified imaging planes or distributed throughout the entire muscle volume. However, planar seeding methods that rely on too few slices may lead to incomplete muscle coverage^19,20^. These voxel-based seeding approaches do not require the aponeurosis to be defined; and if it is defined, the propagated tracts do not necessarily intersect with it. Therefore, this method only supports calculation of the angle α. However, a voxel-based approach may facilitate the analysis of multi-muscle datasets and would provide sufficient information for many musculoskeletal modeling applications.

The goals of this study were to implement and evaluate 1) an updated rounding procedure for aponeurosis-based tract seeding to improve the quantitative accuracy of the tract length estimates and 2) several options for voxel-based seeding. The voxel-seeding options included regularly spaced voxels throughout muscle volume; a variable density seeding approach intended to improve the spatial uniformity of fiber tract density; and an edge-seeding method in which seed points were placed one voxel internal to the muscle’s boundaries. We first implemented and evaluated these approaches using simulated datasets. The tractography outcomes were evaluated in terms of their agreement with the ground truth fiber orientations at the voxel level, the correct termination of tracts at the muscle’s outer boundary, and noise sensitivity. We subsequently demonstrated the methods’ implementation in an example human dataset.

## 2. Methods

### 2.1 Simulations

#### 2.1.1 Definition of Model Tissue and Noise-free Images

The simulation strategy was adapted from previously described methods^10,14,21^. A composite tissue, including two approximately symmetric muscular halves placed about a central aponeurosis, was modeled (Figure 1). Along the long axis of the tissue (taken as the Z direction), 37 slices of tissue were defined with an in-plane resolution of 1×1 mm^2^ and 6-mm thickness. Detailed procedures for modeling fiber orientations are given in the Supplement. Briefly, the fibers projected outward from points along the aponeurosis to corresponding points on the muscle’s outer boundary (defining the azimuthal angle, □; Figure 1A-C). The fibers had angles of elevation (θ) of 66.4-82.0□ (mean, 72.4□; Figure 1D-F). The θ increased as a function of ascending slice number and as a function of distance from the aponeurosis. In each voxel, the unit-length fiber orientation vector was taken as ε_1_, the first eigenvector of the diffusion tensor **D**. ε_3_ was calculated as the cross product of ε_1_ and a unit vector in the +Z direction and ε_2_ was calculated as the cross product of ε_1_ and ε_3_. For each muscle voxel, **D** was calculated as:

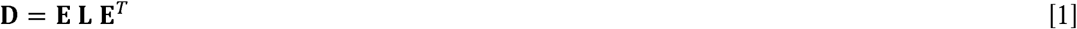

**Figure 1.**
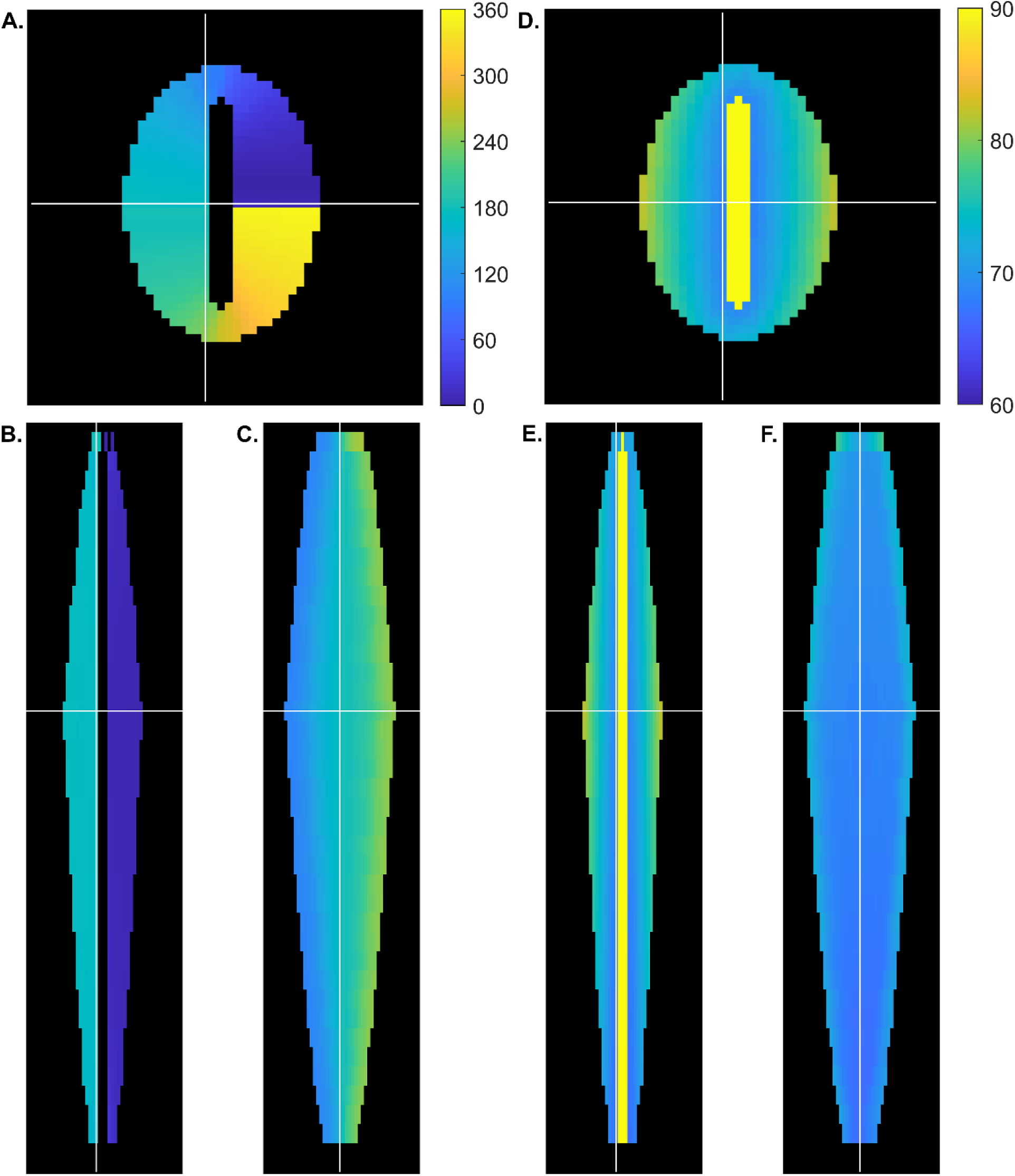
Composite tissue design and muscle fiber orientations in the simulated muscle. Panels A-C show angles of azimuth (□) as counterclockwise rotations from the +X axis. **A.** Axial view. The elliptical-profiled tissue had a central aponeurosis in which the collagen orientation was perpendicular to the slice plane. White lines show the locations of the coronal (horizontal line) and sagittal (vertical line) views. The color bar gives □ in degrees. **B.** Coronal view. The same color scale as in Panel A. is used. White lines give the locations of the axial (horizontal line) and sagittal (vertical line) views. **C.** Sagittal view. The angle of decreased as a function of increasing slice number. The same color scale as in Panel A. is used. White lines give the locations of the axial (horizontal line) and coronal (vertical line) views. **D-F.** Same as A-C, except that the angle of elevation above the slice plane (θ) is shown. **D.** Axial view. The color scale indicates the θ in degrees. θ increased as a function of within-slice distance from the aponeurosis. **E.** Coronal view. The same color scale as **D.** is shown. θ increased as a function of increasing slice number. **F.** Sagittal view. The same color scale as **D.** is shown. θ increased as a function of increasing slice number.

With **E** the eigenvector matrix, **L** a diagonal matrix of eigenvalues, and the superscript *T* indicating matrix transposition. The eigenvalues of **L** were assumed to be 2.1, 1.6, and 1.4 ×10^−5^ cm^2^/s.

Noise-free images were calculated for a matrix size of 50×50 and 40 slices, with one slice proximal to the muscle and two slices distal to the muscle being padded with zeros. Diffusion-encoding used a non-diffusion weighted image and 12 diffusion-weighted images, defined using a diffusion-weighting (b-) value of 475 cm^2^/s and diffusion-encoding directions defined by the minimum energy arrangement of points on a sphere^22^. The signals were calculated as:

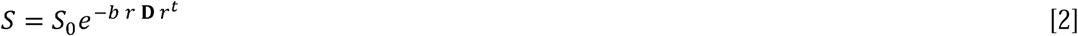

With *S* the diffusion-weighted signal, *S_0_* the non-diffusion-weighted signal (=1), and *r* a three-element vector specifying each diffusion-encoding direction.

#### 2.1.2 Evaluation of Aponeurosis Seeding Changes in the Noise-free Condition

As noted, a seeding mesh dilation process can be used to overcome errors caused by digitization steps, partial volume artifacts, and rounding conventions; but it may underestimate the true fascicle length. To obviate this dilation process, an updated method for initiating fiber tracts from the points defining a seeding mesh was implemented and evaluated. Specifically, we introduce the following procedure to ensure that any seed point placed within one voxel of the muscle-aponeurosis boundary can be used to initiate a fiber tract, while still beginning at the preferred seed point:

1. Form masks defining the internal aponeurosis boundaries and the muscle boundaries (the latter excluding the internal aponeurosis).
2. Define an aponeurosis seeding mesh at the muscle-aponeurosis boundary. Define each vertex in this mesh as a seed point and find its row, column, and slice coordinates.
3. Round the slice coordinates using standard rounding conventions; for the row and column coordinates, take both the floor and ceiling.
4. For each of the four combinations of integer row and column coordinates, determine if the point is included within the muscle boundary mask or the aponeurosis boundary mask.
5. For each point included within the muscle boundary mask, determine the distance between the rounded coordinates and the seed point; define the point with the smallest distance as the closest included point.
6. Propagate a tract from the seed point using the diffusion direction found upon lookup using the closest included point.

We performed trials that used an expanded aponeurosis seeding mesh (the lateral mesh points were up to 0.5 mm outside of the true muscle-aponeurosis boundary and therefore within the muscle boundary mask, per standard rounding conventions) and trials that used a restricted aponeurosis seeding mesh (the lateral mesh points were 10 μm inside of the true muscle-aponeurosis boundary and within the aponeurosis boundary mask, per standard rounding conventions). The latter condition was performed strictly to illustrate the error and was not used for quantitative analysis. These conditions were tested first without implementing the seed point rounding algorithm (Conditions APO-1 and APO-2 for the expanded and restricted meshes, respectively) and again using the restricted mesh after such implementation (Condition APO-3). The points defining the restricted and expanded seeding meshes are illustrated in Figure S2.

Fiber tracking was performed using the MuscleDTI_Toolbox, as previously described^14,17,23^ except as updated for the purpose in this work; see the Supplement, Table S1 for a list of function versions used and their modifications from previous publications. Fiber tracts were propagated from the seed points in the +Z direction by Euler integration of ε_1_ at a step-size of one voxel-width. Premature termination due to excessive inter-point angles or fractional anisotropy (FA) values did not occur in the noise-free condition. To smooth the tracts, their row, column, and slice coordinates were fitted to 2^nd^-, 2^nd^, and 3^rd^-order polynomials, respectively, as functions of point number.

The smoothed tracts’ architectural properties were characterized as follows. Tracts with length (L_FT_)>5 mm were analyzed. To promote uniform sampling along the aponeurosis seeding mesh, we first defined redundant fiber tracts as multiple tracts that shared the same seed point (after rounding to the nearest 1 mm^3^). One of the redundant tracts, selected at random, was preserved. For each preserved tract, the α values were calculated in a pointwise manner along the tract by finding the angle formed by the line connecting the fiber-tract point to the seed point and the local tangent to the muscle’s central axis at the level of the seed point. The central axis was determined by fitting 3^rd^-order polynomial functions through the row and column coordinates of each muscle slice’s centroid, as a function of slice number. A vector describing the slice-wise difference in the central axis position, converted to unit length, was taken as the local tangent to the central axis. Curvature (κ) was calculated in a pointwise manner along the tract using the Frenet-Serret formulas, as previously described^21^. The tracts were characterized with their average α and κ values, and the L_FT_ was calculated as the summed Euclidean distance between points. For each architectural property, the whole muscle mean value was calculated.

The fiber orientations within the tracts were evaluated by comparing the θ and □ for each segment of the smoothed fiber tracts to the ground truth data at the corresponding voxel location. The mean angular difference and limits of agreement for the whole muscle were calculated and visualized using Bland-Altman plots^24^, and the type 1,1 intraclass correlation coefficients (ICCs) were calculated between the estimated and ground truth angles. Angles that fell symmetrically about the +X axis (for example, 1° and 359°) were excluded from the analysis of □. Finally, the tract point density in each voxel of the muscle boundary mask was calculated and averaged for each slice.

#### 2.1.3 Implementation and Characterization of Voxel Seeding Methods: Noise-free Condition

Three methods of voxel-based seeding were introduced and evaluated: regularly spaced seeding; variable density seeding; and edge seeding. For regularly spaced seeding (condition VXL-1), seed points were placed in alternating voxels in the row and column directions, within a once-eroded muscle boundary mask. Variable density seeding (condition VXL-2) placed seed points at a variable frequency of an eroded muscle boundary mask. The variable frequency resulted in a seed point in every row and column position of slices that contained ≤100 voxels in the eroded boundary mask; every row and every second column for slices that contained 101-200 voxels in the eroded boundary mask; every row and every third column for slices that contained 201-300 voxels in the eroded boundary mask; etc. to a maximum frequency of one point every row and sixth column for slices that contained 501 or more voxels in the eroded boundary mask. These seed densities were based on preliminary empirical observations and were designed to make the sampling of the muscle by the fiber tract points more spatially uniform. Edge-seeding (condition VXL-3) placed seed points in all voxels that lay along the outer edge of the eroded muscle mask and along the outer edge of the dilated aponeurosis boundary mask. The voxel seeding methods are illustrated in Figures S3-S5.

For each voxel seeding condition, fiber tracts were propagated bidirectionally from the points. Premature termination did not occur in the noise-free condition; however, rounding and tract termination conventions sometimes resulted in tract propagation into the internal aponeurosis region of the muscle. Therefore, up to four innermost points of each fiber tract were removed if they lay inside of the internal aponeurosis boundaries. Tract smoothing was performed as described as in §2.1.2. Tracts with L_FT_>5 mm were analyzed. Redundant tracts, defined as tracts that shared the same innermost fiber-tracking point (rounded to within 1 mm^3^), were treated as described in §2.1.2. Architectural characterization included the L_FT_, mean α, and mean κ, with α calculated by treating the innermost point on the fiber tract as the seed point.

The outcomes were evaluated as described in §2.1.2 and by comparing the architectural properties to those of the noise-free APO-3 condition. To account for the architectural heterogeneity of the simulated muscles, only the tracts whose innermost points had Z positions corresponding to those of the aponeurosis (i.e., the same levels as the seed points of the APO-3 tracts) were included in these analyses.

#### 2.1.4 Comparison of Aponeurosis and Voxel Seeding Using Noisy Data

The performance of four proposed seeding methods (APO-3, VXL-1, VXL-2, and VXL-3) was evaluated in the presence of noisy data. The APO-1 and APO-2 conditions were not further investigated because it will be shown that they underestimated the tract length or resulted in tract initiation failures in the noise-free condition.

The following procedures were performed at SNR levels of 39 and 54. For each of 1000 independent trials, Gaussian-distributed noise was added to the noise-free images, as previously described^14^. The noisy images were used to estimate **D** using a weighted least squares method, followed by singular value decomposition to obtain estimates of **E** and **L**. Fiber tracts were propagated and analyzed using the procedures described above, except that premature tract termination could occur if either the FA value fell outside range of 0.1-0.4 or the angle between the propagated segment and the second segment preceding the propagated segment exceeded 30□ for two consecutive points. The fiber tracts were smoothed as described as in §2.1.2, selected for L_FT_>5 mm, filtered for redundancy at the innermost fiber tract point (voxel seeding) or seed point (aponeurosis seeding), and characterized with their L_FT_, average α, and average κ. These data were averaged across the entire muscle. At the end of the 1000 trials, the average value of each parameter was calculated, and actual 95% confidence intervals were formed from the data points at the 2.5^th^ and 97.5^th^ percentiles.

The outcomes were evaluated by comparison of the architectural properties to those of the noise-free APO-3 condition. For these comparisons, only the tracts whose innermost points had Z positions corresponding to those of the aponeurosis (i.e., the same levels as the seed points of the APO-3 tracts) were included.

### 2.2 Demonstration in Human Data

The performance of the four candidate seeding methods (APO-3, VXL-1, VXL-2, and VXL-3) was also demonstrated with *in vivo* data. These procedures were approved by our local Institutional Review Board. A 21-year-old female provided informed consent to participate. MRI used a Siemens Vida, XA30 3T MRI scanner. The participant was positioned supine on a patient bed containing built-in spine elements. An 18-element array coil was placed over the anterior surface of the leg and supported by a frame running along the leg’s medial and lateral surfaces. The heel and thigh were supported to raise them above the patient bed, allowing the posterior leg muscles to be freely suspended. The body coil was used for transmission; receive coil selection used the manufacturer’s “Auto Coil Select” function.

Structural imaging data were acquired using Siemens’ product implementation of quantitative fat-water imaging (Volume Interpolated Breath-hold Examination, VIBE). Three-dimensional (3D) Fast Low-Angle SHot gradient echo images were acquired in two stacks with matrix size, 160×160 acquired (320×320 reconstructed), field of view (FOV), 200×200 mm^2^; 64 slices with slice thickness (ST), 3 mm and inter-slice gap, 0 mm; repetition time (TR) = 9.23 ms; echo time (TEs), [1.37, 2.60,…,7.52] ms; flip angle, 4□; and number of excitations (N_EX_), 1. DT-MRI data were also acquired in two stacks. A REadout Segmentation Of Long Variable Echo-trains (RESOLVE) sequence was used with matrix size, 72×72 acquired (144×144 reconstructed), FOV, 200×200 mm^2^; 32 slices per stack with ST, 6 mm and inter-slice gap, 0 mm; TR = 3700 ms; TE, 46.18 ms; b-value, 0/450 s/mm^2^ with 10 directions; N_EX_ for b=0/450, 4/2; 7 readout segments; and “strong” (fat saturation and gradient reversal) fat suppression. The DT-MRI slice offsets were defined to overlap with contiguous pairs of VIBE slices. A four-slice overlap between the stacks was used for the VIBE dataset and a two-slice overlap was used for the RESOLVE dataset. The shim volume was defined to cover the lower-leg muscles in each stack, while minimizing fat and air coverage. The full width at half maximum (FWHM) peak height was measured prior to the RESOLVE scans. A FWHM of 43.8 Hz was measured prior to the proximal RESOLVE image stack, and a FWHM of 64.0 Hz was measured prior to the distal RESOLVE image stack.

The VIBE data were processed using the manufacturer’s algorithms to produce maps of the fat and water signal distributions. For both VIBE and RESOLVE, an offset was observed between the corresponding slices in the overlapping regions of the proximal and distal image stacks. The artifact was corrected by registering the first stacks of the VIBE and RESOLVE datasets to their respective second stack with a 2D rigid body translation using MATLAB’s *imwarp( )* function. The inter-stack 2D translations were defined using the *imregtform( )* function, with one transformation obtained from each pair of overlapping slices, and a final transformation defined for each dataset by averaging the 2D transformations of the overlapping slices of each respective VIBE and RESOLVE stack pair. The two stacks of each dataset were concatenated after eliminating the overlapping slices on the distal image stack. Finally, the VIBE dataset was resized to match the RESOLVE dataset, and the RESOLVE dataset was registered to the resized VIBE dataset with a 2D rigid body translation transformation. The 2D translation applied to the RESOLVE dataset was defined by averaging the 2D transformations obtained after individually registering the central 50 slices of both datasets using the *imregtform()* function.

The registered RESOLVE dataset was denoised using anisotropic smoothing^25,26^ at a level of 5%^27^. The lower leg’s diffusion tensor field was computed from the denoised dataset using a weighted linear least squares method. Fiber tracts were generated in the tibialis anterior (TA) muscle using the APO-3, VXL-1, VXL-2, and VXL-3 methods. The TA muscle boundary mask was manually segmented and used in all methods. For APO-3, the aponeurosis was segmented and formed into a 35×45 mesh using the *define_roi( )* function in the MuscleDTI_Toolbox^17^. Voxel-based seeding was implemented as described above. Fiber tracts were propagated using Euler integration at a step-size of one voxel-width. Tract termination occurred at the muscle boundary or either the FA value fell outside range of 0.05-0.40 or the angle between the propagated segment and the second segment preceding the propagated segment exceeded 30□ for two consecutive points. The tracts were selected for L_FT_>10 mm and <104 mm, filtered for redundancy, and characterized with their L_FT_, average α, and average κ. For APO-3, γ was calculated as the pennation angle in the MuscleDTI_Toolbox^17^, and β was calculated as (α – γ). The whole muscle average values were calculated. The outcomes of the voxel-based seeding were evaluated by comparison of the architectural properties to those of the APO-3 condition. For these comparisons, only the tracts whose innermost points had Z positions corresponding to those of the aponeurosis (i.e., the same levels as the seed points of the APO-3 tracts) were included.

## 3. Results

### 3.1 Simulations

#### 3.1.1 Aponeurosis Seeding Changes in the Noise-free Condition

Figure 2A shows fiber tracts generated in condition APO-1, in which the expanded aponeurosis seeding mesh and current rounding method were used. When the restricted aponeurosis mesh and current rounding method were used (condition APO-2), tract propagation on the lateral portions of the aponeurosis mesh failed (Figure 2B). Use of the proposed seed point rounding method with the restricted mesh (condition APO-3) resulted in tracts that extended from the aponeurosis seeding mesh to the outer boundaries; see Figure 2C.

**Figure 2.**
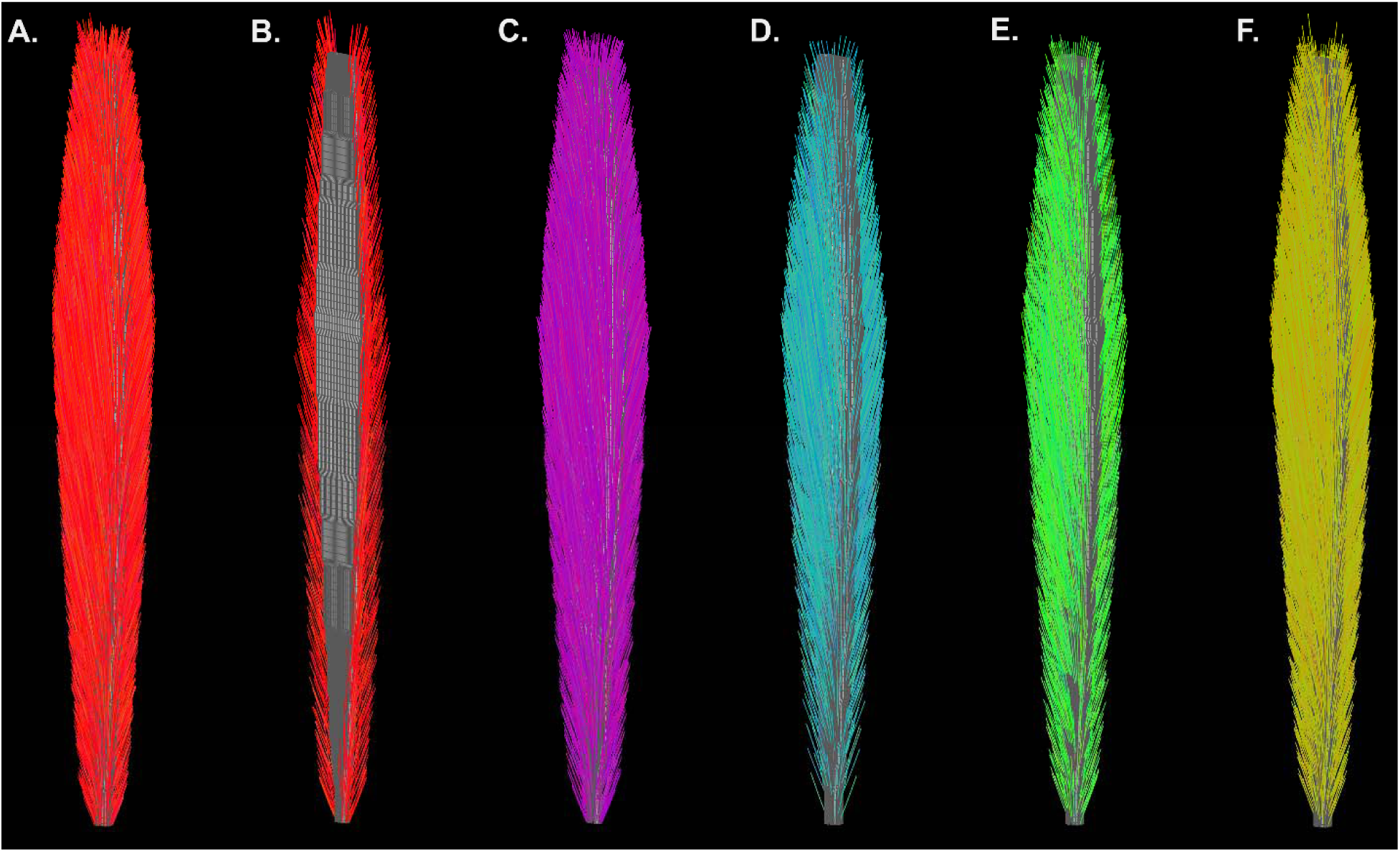
Fiber tracts generated under noise-free conditions (simulated muscle). **A.** Fiber tracts generated using the current tracking methods, with an expanded aponeurosis seeding mesh whose lateral mesh column coordinates fell 0.2-0.5 pixels (0.2-0.5 mm) outside the rounding limits that allow fiber tract initiation (APO-1). **B.** Fiber tracts generated using the current tracking methods, with a restricted aponeurosis seeding mesh whose lateral mesh column coordinates fell 0.01 pixels (10 μm) inside the rounding limits that allow fiber tract initiation (APO-2). **C.** Aponeurosis-seeded fiber tracts generated using the restricted mesh and the updated seed point rounding procedure (APO-3). **D.** Regularly spaced, voxel-seeded fiber tracts (VXL-1). **E.** Variable density, voxel-seeded fiber tracts (VXL-2). **F.** Edge voxel-seeded fiber tracts (VXL-3).

The architectural estimates in the APO-1 and APO-2 seeding conditions are shown in Table 1. By definition, tracts generated from the restricted mesh started within 10 um of the muscle-aponeurosis boundary, while those generated from the expanded mesh started up to 0.5 mm outside of this boundary. All tracts ended at the outer muscle boundary. Figure 3 shows the agreement between the θ and □ angles known from ground truth and those estimated from the APO-3 fiber tracts. The type 1,1 ICC and its 95% confidence interval for the θ and □ angles were 0.949 [0.948 0.951] and 0.990 [0.990 0.991], respectively. The mean difference and [limits of agreement] for the θ and □ angle differences were 0.9 [0.0 1.5] and 0.2 [-5.8 6.3], respectively, with the unrounded lower limit of agreement for θ being negative. However, the pattern of the differences in the θ angle suggests that the tract-based estimates slightly overestimated the known θ at low values. Also, the variability in the mean difference in □ reached local minima for values of about 180 and 360□; and there was a relative paucity of data points near 90 and 180°. The average κ was 2.2% greater for the APO-1 condition than for the APO-3 condition.

**Figure 3.**
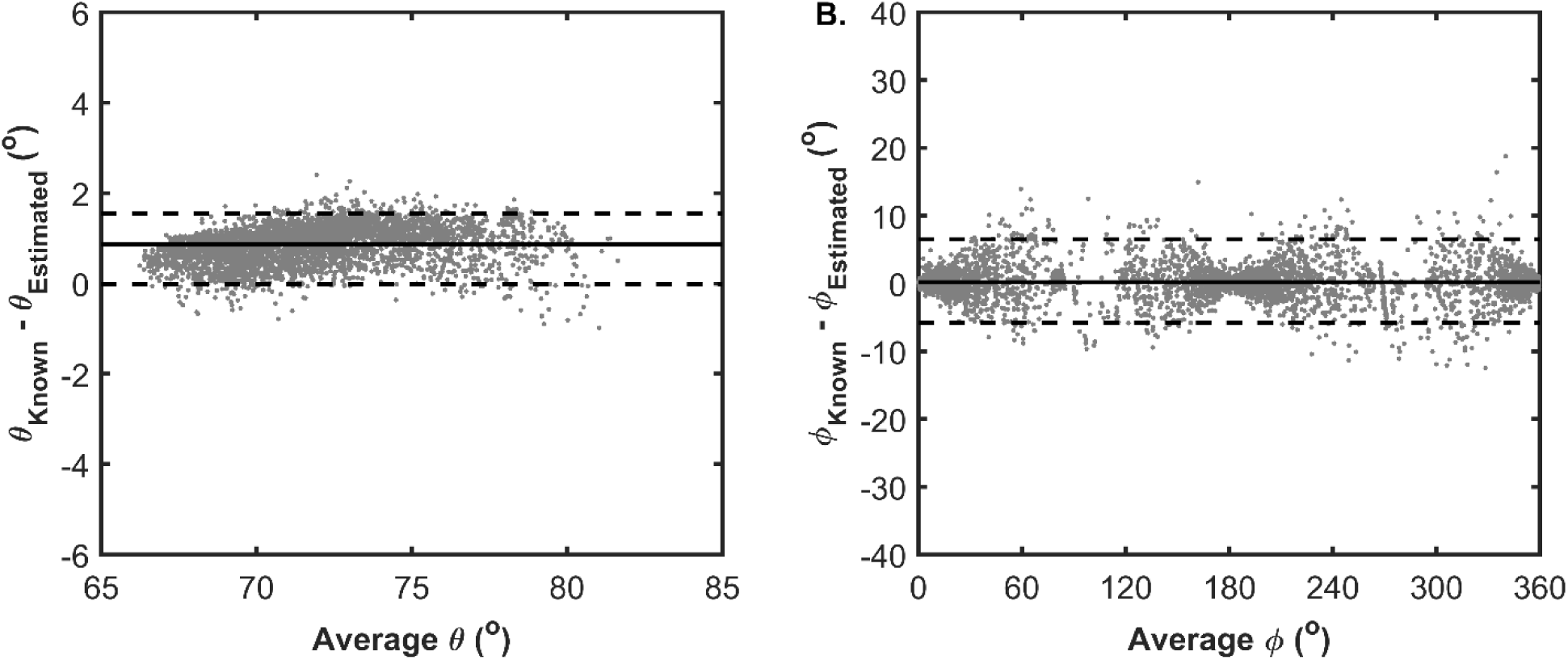
Bland-Altman plots of the agreement between of θ and □ for the fiber tract segment angles and ground truth (simulated muscle). **A.** The mean pairwise difference (ground truth – tract segment-based estimate) in θ is plotted as a function of the mean pairwise value of θ. The actual 95% confidence interval (i.e., that calculated from the 2.5^th^ and 97.5^th^ percentiles of the distribution of difference scores) is shown. included zero. For clarity, only every 10^th^ data point is shown. **B.** The mean pairwise difference (ground truth – tract segment-based estimate) in □ is plotted as a function of the mean pairwise value of □. The 95% confidence interval included zero. For clarity, only every 10^th^ data point is shown.

**Table 1.** Architectural Estimates in Each SNR Condition. The mean and [actual 95% confidence intervals] are given for the SNR=39 and 54 conditions.

| Seed Method | SNR | $L_{FT}$ (mm) | $\alpha$ ($^{\circ}$) | $\kappa$ ($m^{-1}$) |
| --- | --- | --- | --- | --- |
| APO-1 | Noise-free | 19.7 | 20.0 | 6.5 |
| APO-3 | Noise-free | 21.2 | 20.2 | 6.4 |
|  | 39 | 21.6 [21.3 21.9] | 20.0 [19.8 20.2] | 12.7 [12.3 13.1] |
|  | 54 | 21.7 [21.5 21.8] | 20.0 [19.9 20.2] | 10.2 [9.0 10.5] |
| VXL-1 | Noise-free | 23.4 | 19.5 | 5.5 |
|  | 39 | 24.6 [24.1 25.0] | 19.1 [18.8 19.3] | 10.6 [10.2 11.0] |
|  | 54 | 24.4 [24.1 24.7] | 19.2 [19.0 19.4] | 8.6 [8.3 8.9] |
| VXL-2 | Noise-free | 25.4 | 19.2 | 5.6 |
|  | 39 | 26.3 [25.9 26.8] | 18.8 [18.6 19.0] | 10.0 [9.6 10.4] |
|  | 54 | 26.2 [25.9 26.5] | 19.0 [18.8 19.1] | 8.2 [8.0 8.5] |
| VXL-3 | Noise-free | 22.3 | 19.5 | 5.6 |
|  | 39 | 23.1 [22.7 23.4] | 19.2 [19.0 19.4] | 11.3 [10.9 11.6] |
|  | 54 | 22.9 [22.7 23.2] | 19.3 [19.2 19.4] | 9.1 [8.8 9.3] |

Figure 4A shows the slice-wise average number of fiber-tracking points per voxel. For slices having at least two proximal and distal neighboring slices, the average number of fiber tract points/voxel in each slice was 6.6 (standard deviation, 0.8; coefficient of variation, 12.4%).

**Figure 4.**
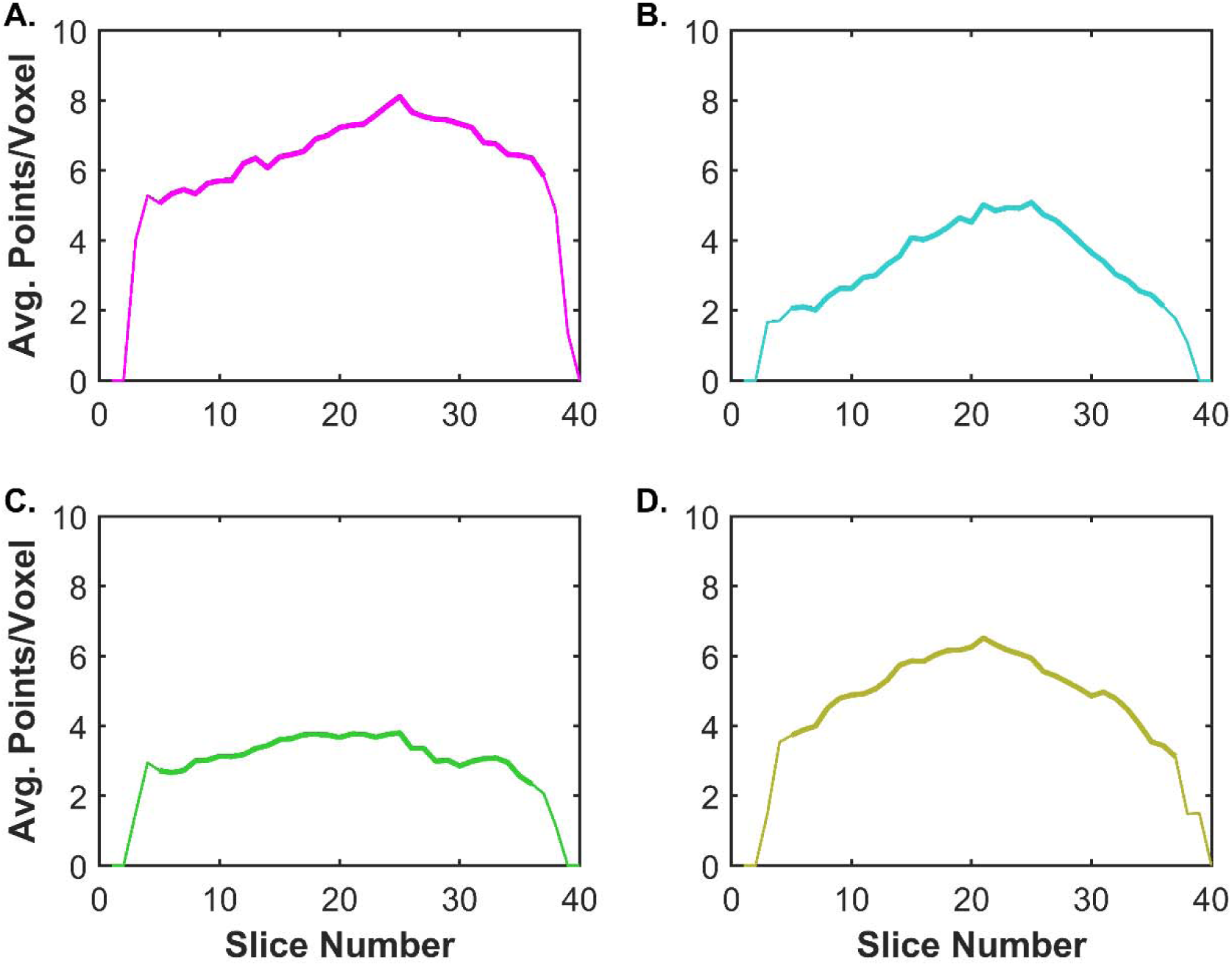
Uniformity of spatial sampling of the muscle by the fiber tracts (simulated muscle). Uniformity is expressed as the average number of fiber tract points per voxel in each slice. Descriptive statistics were calculated for slices with at least two proximal and distal neighbors, as represented by the bold lines. **A.** APO-3 condition, with a mean of 6.6 points/voxel and a coefficient of variation (CV) of 12.4% across slices. **B.** VXL-1 condition, with a mean of 3.6 points/voxel and a CV of 28.4%. **C.** VXL-2 condition, with a mean of 3.2 points/voxel and a CV of 12.9%. **D.** VXL-3 condition, with a mean of 5.1 points/voxel and a CV of 18.5%.

#### 3.1.2 Implementation and Characterization of Voxel Seeding Methods: Noise-free Condition

Figures 2D-F show the fiber tracts generated using regularly spaced voxel-seeding, variable density seeding, and edge seeding, respectively. The architectural estimates are provided in Table 1. For L_FT_, the values under the VXL-1, VXL-2, and VXL-3 conditions were greater than under the APO-3 condition. For all voxel seeding conditions, the whole-muscle average α and κ estimates were lower than for APO-3. Figures 4B-D show the uniformity of spatial sampling of the muscle by the fiber tracts for the VXL-1, VXL-2, and VXL-3 conditions. The average and [standard deviation, coefficient of variation] numbers of fiber tract points/voxel in each slice were 3.6 ([1.0, 28.4%], 3.2 [0.4, 12.9%], and 5.1 [0.9, 18.5%] for VXL-1, VXL-2, and VXL-3, respectively.

#### 3.1.3 Comparison of Aponeurosis and Voxel Seeding Using Noisy Data

Figure 5 shows example fiber tracts generated using images with SNR=39 under the APO-3, VXL-1, VXL-2, and VXL-3 conditions. The architectural estimates for the SNR=39 and 54 conditions are provided in Table 1. In general, the addition of noise to the images tended to cause L_FT_ to be slightly overestimated and α to be slightly underestimated, with no systematic effect of SNR level. However, κ was consistently overestimated in an SNR-dependent manner.

**Figure 5.**
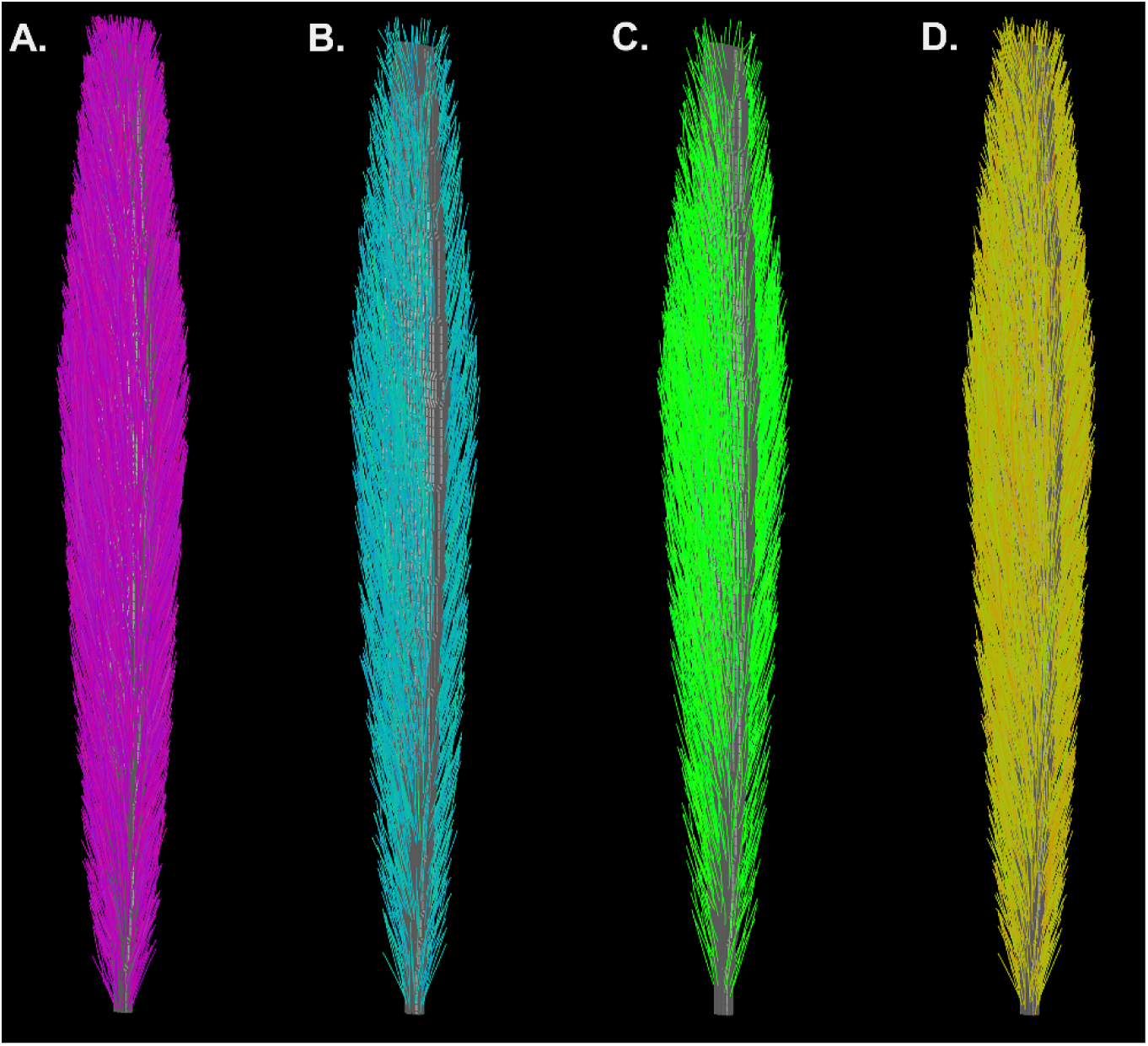
Example fiber tracts generated in the SNR=39 condition (simulated muscle). **A.** APO-3 condition (restricted aponeurosis seeding mesh with updated rounding procedure). **B.** VXL-2 (variable density seeding) condition. **C.** VXL-3 (edge seeding) condition.

### 3.2 Demonstration in Human Data

Figure 6 shows fiber-tracts generated from human imaging data. The architectural estimates are provided in Table 2. In general, the L_FT_ values for the VXL conditions were slightly lower than for the APO-3 condition, and the α angles were slightly larger than for APO-3. The β and γ angles are also reported for APO-3. There was not a systematic effect of voxel-seeding on κ, in comparison to the APO-3 condition. Figure 7 shows the slice-wise average number of fiber-tract points per voxel for each seeding method. The average and [standard deviation, coefficient of variation] numbers of fiber-tract points/voxel in each slice were 8.0 [1.3, 16.2%], 5.9 [1.4, 23.6%], 7.4 [1.4, 19.2%], and 8.5 (2.4, 28.9%) for APO-3, VXL-1, VXL-2, and VXL-3, respectively.

**Figure 6.**
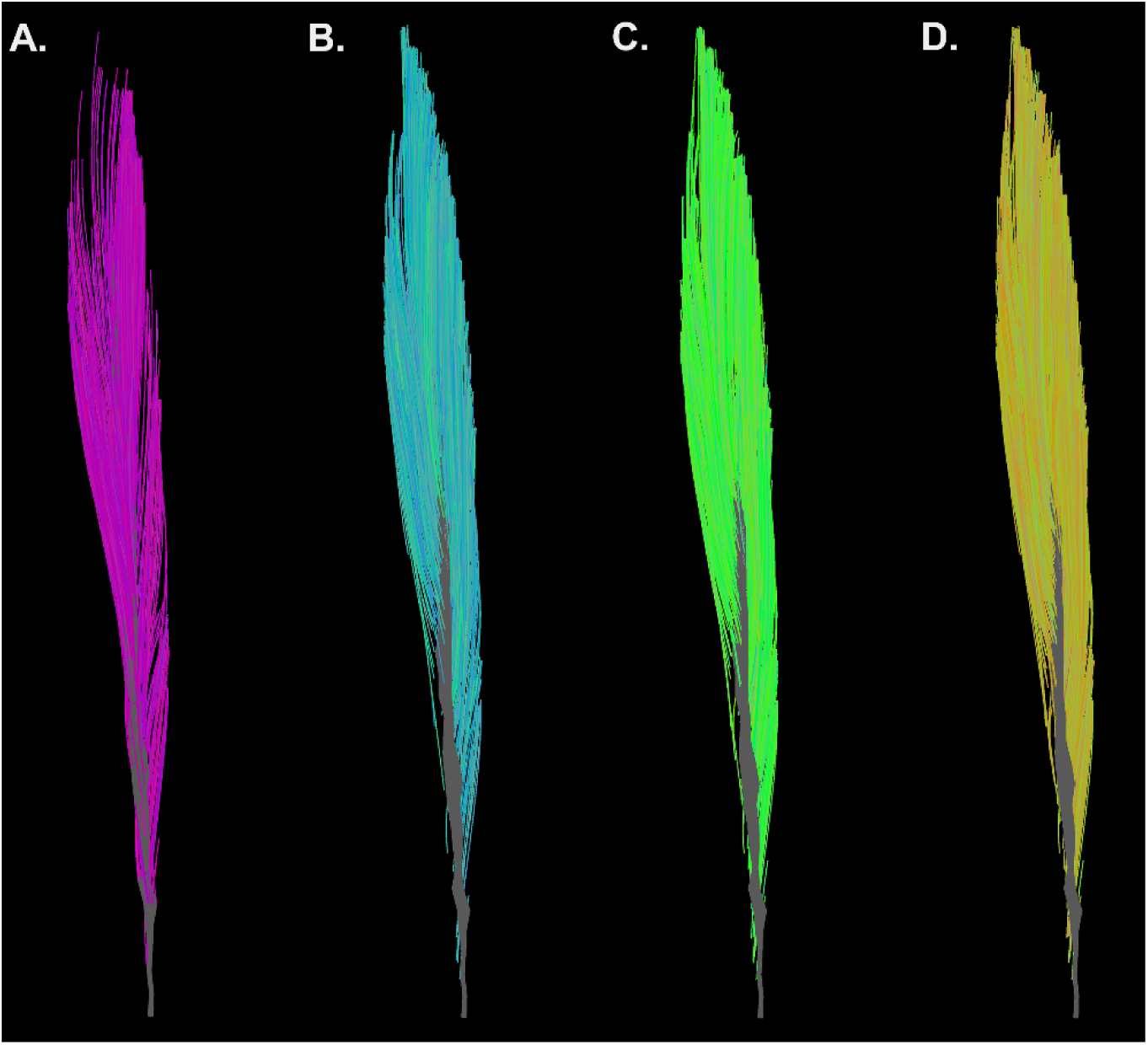
Example fiber tracts (*in vivo* data). **A.** APO-3 condition (restricted aponeurosis seeding mesh with updated rounding procedure). **B.** VXL-1 (every 2^nd^ voxel in the row and column directions). **C.** VXL-2 (variable density seeding) condition. **D.** VXL-3 (eroded edge seeding) condition.

**Figure 7.**
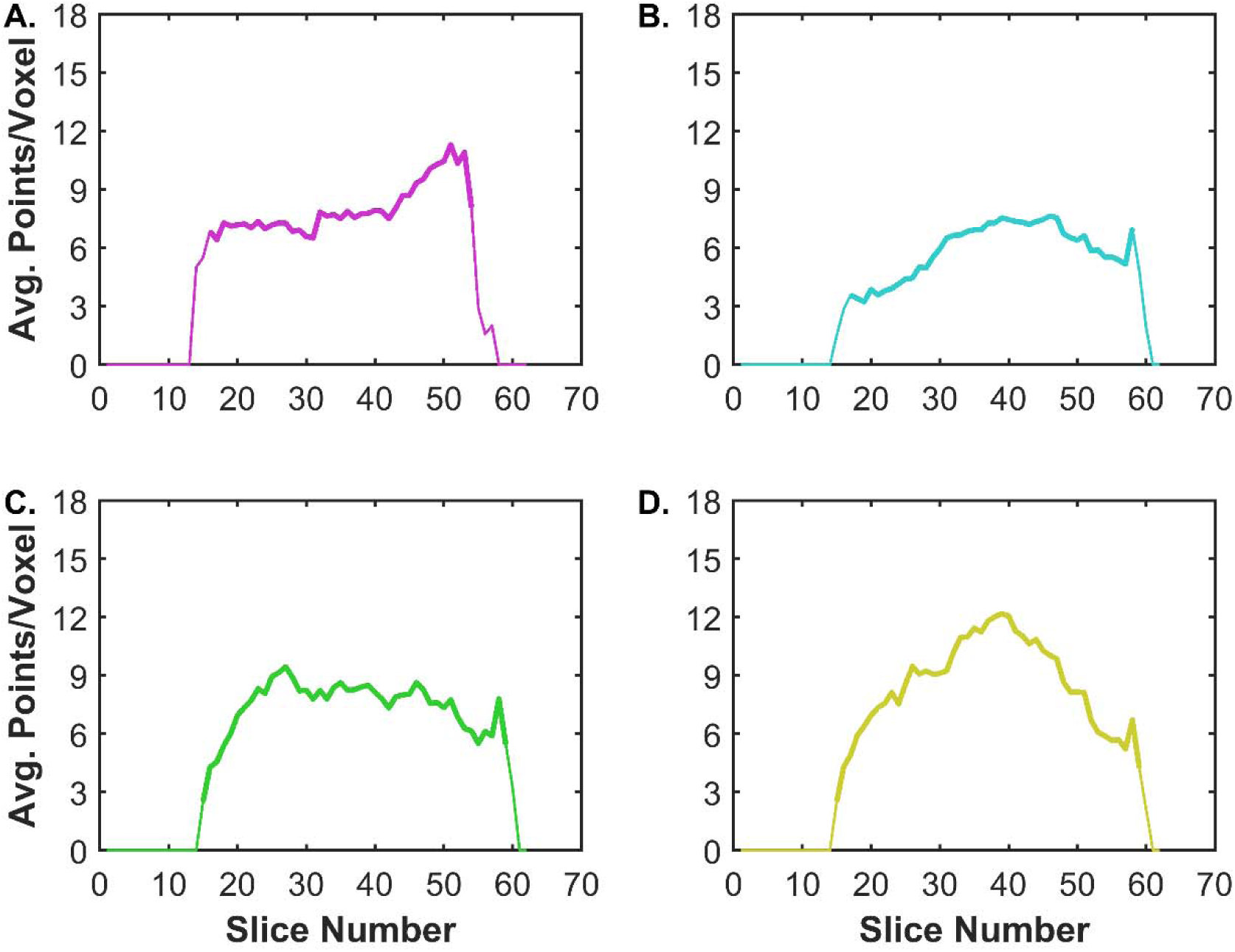
Uniformity of spatial sampling of the muscle by the fiber tracts (*in vivo* data). Uniformity is expressed as the average number of fiber tract points per voxel in each slice. Descriptive statistics were calculated for slices with at least two proximal and distal neighbors, as represented by the bold lines. **A.** APO-3 condition, with a mean of 8.0 points/voxel and a coefficient of variation (CV) of 16.2% across slices. **B.** VXL-1 condition, with a mean of 5.9 points/voxel and a CV of 23.6%. **C.** VXL-2 condition, with a mean of 17.4 points/voxel and a CV of 19.2%. **D.** VXL-3 condition, with a mean of 8.5 points/voxel and a CV of 28.9%.

**Table 2.** Architectural Estimates in Human Muscle. The mean values are given for the APO-3, VXL-1, VXL-2, and VXL-3 conditions. α is the angle made by the fiber tracts and the central axis of the muscle. β and γ, respectively, are the angles made by the aponeurosis and the central axis of the muscle and by the fiber tracts and the aponeurosis.

| Seed Method | $L_{FT}$ (mm) | Angles ($^{\circ}$) | | | $\kappa$ ($m^{-1}$) |
| --- | --- | --- | --- | --- | --- |
| | | $\alpha$ | $\beta$ | $\gamma$ | |
| APO-3 | 73.3 | 9.2 | 2.5 | 6.7 | 4.2 |
| VXL-1 | 72.9 | 9.4 |  |  | 4.1 |
| VXL-2 | 72.4 | 10.0 |  |  | 4.4 |
| VXL-3 | 70.7 | 10.0 |  |  | 4.3 |

## 4. Discussion

### 4.1 Modeling of Muscle Architecture for DT-MRI Tractography Simulations

The present simulation approach is based on previous uses of simulated muscle architectures for code validation and understanding the effects of image acquisition and tractography settings on the accuracy and precision of DT-MRI tractography-derived architecture estimates^10,11,14,17,21^. We further expand on these approaches by using two mechanisms to create architectural heterogeneity. The first mechanism was the muscle’s overall morphology: the muscle’s minor and major radii varied as a function of slice number, creating heterogeneity in muscle fiber length. In addition, fiber orientation was varied: the angles of azimuth were distributed in an approximately radial pattern and the angles of elevation increased with both ascending slice number and in-plane distance from the center of the muscle. Importantly, these properties of the simulated muscle model the morphological variations and the variations in fiber orientation present in many human muscles. For example, in the vastus medialis muscle, the pennation angle increases from 5□ to 50□ (Ref.^28^) and the muscle thickens it proceeds from origin to insertion. Moreover, Blemker *et al.* have shown that architectural heterogeneity can induce heterogeneous patterns of strain development during contraction^6^, and recent finite element models of muscle that incorporate heterogeneous fiber architecture have been used to predict fascicle and tissue behavior in passive and active muscle lengthening^29^. The existence and functional impact of architectural heterogeneity therefore illustrates the importance of using heterogeneous architectures for applications such as this, as well as developing seeding methods that promote uniform spatial sampling by the fiber-tracts.

### 4.2 Aponeurosis Seeding Strategies

Aponeurosis seeding is an example of tract propagation from functionally relevant seed points, much like using functional MRI-derived regions of interest as seed points for white matter tractography. For skeletal muscle tractography, the functional relevance of aponeurosis seeding is that the aponeurosis is the structure into which most of the muscle fascicles insert and through which most of the fibers’ force is transmitted. As introduced here, aponeurosis-seeding also allows the estimation of additional functionally relevant structure properties, including its size and orientation with respect to the muscle’s fibers and mechanical line of action.

White matter tractography based on seed points derived from functional MRI-defined regions of interest can encounter errors due to partial volume artifacts between white and gray matter^30^. Likewise, partial volume artifacts between muscle and aponeurosis can induce ε_1_ estimation errors in skeletal muscle tractography. This error can be avoided by placing the seeding mesh outside of the muscle-aponeurosis boundary. For example, the use of the expanded seeding mesh – which was placed up to 0.5 mm outside of this boundary – allowed tract initiation to occur on a locally correct trajectory even when conventional rounding conventions were applied. However, for the angles of elevation that we modeled, using the expanded seeding mesh also led to a ∼1.5 mm underestimation of the fiber tract length. The updated seed point rounding algorithm corrected this error, improving the estimate of tract length.

The APO-3 condition had acceptable levels of accuracy and precision in its fiber orientation estimates. The mean difference for elevation angle had a 95% confidence interval that included zero; and the value-dependent bias that did exist was small (<1□) and would therefore not be expanded to significantly impact muscle architecture-based predictions of force production (which vary with the cosine of the pennation angle). The estimate of the azimuthal component of ε_1_ was less biased than the estimate of θ, but also less precise; but as this is a minor component to ε_1_ in most muscles, the practical impact of the relatively lower precision is expected to be small. However, a persistent source of potential error in aponeurosis seeding is that the fiber-tract point density per voxel varied in a manner that apparently corresponds to the muscle and aponeurosis cross-sectional area in the slice. This was observed in both the simulated and human muscles. The impact of this error is likely to depend on the existence and nature of any architecture heterogeneity within the muscle. Other errors were due to noise: at SNR levels of 39 and 54, the tract lengths were slightly overestimated, and the curvature levels were overestimated in an SNR-dependent manner, consistent with previous findings^14,21^.

### 4.3 Voxel Seeding Approaches

Another disadvantage to aponeurosis-seeding is the time-consuming nature of region definition. This is especially true for manual definition approaches, but even automated selection methods may require manual inspection. The voxel-based seeding approaches implemented here obviate these manual steps. Although these approaches do not allow the aponeurosis orientation with respect to the fibers of the mechanical line of action to be estimated, the information that they do provide may continue to be useful in many musculoskeletal modeling applications.

In simulated muscle, in both noise-free and noisy images, the voxel-seeding methods resulted in estimates of tract length (higher), pennation angle (lower), and curvature (lower) that differed from those of the APO-3 condition. Only the curvature estimates demonstrated a clear SNR dependence. For human muscle, however, any differences from the APO-3 condition were both less systematic and lower in magnitude.

The VXL-1 method was implemented here at two voxel-spacing in both the row and column directions. This method is similar to whole volume seeding methods that are used in white matter tractography. Like those approaches, they ensure whole-organ coverage, which might otherwise not occur if excessive FA or high inter-point angle criteria terminate a propagating tract before it reaches a muscle boundary. But also as in white matter tractography – in which large white matter bundles may be overrepresented by a biased distribution of seed points^31^ – VXL-1 seeding resulted in a non-uniform tract density across the slices. Based on the empirical observations in Figure 4B, a simple scheme was developed to vary the seed point density in a slice-specific manner, as a function of the size of the muscle boundary mask (VXL-2). In both simulated and human muscle, this reduced the variability in tract point density across the muscle slices. As implemented here, VXL-2 is similar to seed point definition using planar regions of interest. As noted by Budzik *et al*.^20^ and as illustrated in the review by Damon *et al*.^19^, it is necessary to define multiple planes when seeding based on planar regions of interest, because not all of the fibers in a pennate muscle will pass through a single anatomical plane. By customizing the seed point spacing, the VXL-2 method expands on the multi-planar approaches while also tending to equalize the point density. The VXL-3 (edge-) seeding method provides a simple alternative approach to implement, but also suffers from highly variable tract point densities.

### 4.4 Potential Applications of Updated Aponeurosis and Voxel Seeding Approaches

The APO-3 and VXL seeding methods may also enable additional applications or overcome tractography challenges in diseased muscle. For example, a potential application for the updated aponeurosis seeding method relates to the recent introduction of Laplacian flow simulation to recreate muscle architecture^32^ and support finite element modeling of muscle mechanics^29^. These flow simulations can be validated with DTI tractography data; however, they are sensitive to the aponeurosis definition^33^. Improved aponeurosis seeding methods such as APO-3 can provide more robust methods to validate Laplacian flow simulations or other methods for recreating muscle architecture and modeling muscle function. A potential use of the voxel-based seeding methods is that they may be more robust to early termination or discontinuities of fiber tracts caused by compositional heterogeneity in muscle disease, such as fibrosis^34^, fat infiltration^35^, or muscle fiber branching^36^. While aponeurosis-seeding based tracts could terminate mid-muscle due to partial volume artifacts caused by fibrosis/fat replacement, voxel-based seeding methods are more likely to uniformly sample the muscle regardless of fiber tract discontinuities, as fiber tracts will be initiated from different areas of the muscle.

### 4.5 Conclusions

We have implemented and evaluated several updated fiber-tract seeding methods and tract quantification capabilities. Updated aponeurosis seeding procedures allow more accurate and robust tract propagation from seed points, as well as a broader range of structural quantification. The voxel-based seeding methods implemented here produced quantification outcomes that were closely approximated or well matched to the updated aponeurosis seeding method and are expected to accelerate high throughput analyses of multi-muscle datasets. Continued evaluation of these methods, in a wider range of muscle architectures, is warranted.

## Alphabetized List of Abbreviations and Symbols

APO-1: Original aponeurosis seeding scheme, with expanded mesh coordinates
APO-2: Original aponeurosis seeding scheme, with restricted mesh coordinates
APO-3: Updated aponeurosis seeding scheme
b-value: Diffusion-weighting value
D: Diffusion tensor matrix
DT: Diffusion tensor
DTI: Diffusion tensor imaging
E: Eigenvector matrix
FA: Fractional anisotropy
FOV: Field of view
FWHM: Full width at half-maximum peak height
ICC: Intraclass correlation coefficient
L: Eigenvalue matrix
L_FT_: Fiber tract length
MRI: Magnetic Resonance Imaging
N_EX_: N umber of excitations for signal averaging
r: Diffusion-encoding direction vector
RESOLVE: REadout Segmentation Of Long Variable Echo-trains
S: Observed signal
S_0_: Equilibrium signal intensity
SNR: Signal-to-noise ratio
ST: Slice thickness
TE: Echo time
TR: Repetition time
VIBE: Volume Interpolated Breath-hold Examination
VXL-1: Voxel-based, regularly spaced seeding scheme
VXL-2: Voxel-based, variable density seeding scheme
VXL-3: Voxel-based, edge seeding scheme
α: Angle formed by the muscle’s fibers and its mechanical line of action
β: Angle formed by the muscle’s aponeurosis and its mechanical line of action
γ: Pennation angle
ε_N_: Eigenvector of the diffusion tensor, with N = 1, 2, or 3
θ: Angle of elevation
κ: Curvature
λ_N_: Eigenvalue of the diffusion tensor, with N = 1, 2, or 3
□: Angle of azimuth

## 5. Acknowledgements

The authors thank Profs. Zhaohua Ding, PhD; Melissa Hooijmans, PhD; and Mariana Kersh, PhD for helpful comments.

The authors acknowledge grant support from NIH/NIAMS 1 R01 AR073831.

## Notes

### Competing Interest Statement

The authors have declared no competing interest.

