## Supplement for "A Comparison of Skeletal Muscle Diffusion Tensor Imaging Tractography Seeding Methods"

**S1. Detailed Methods: Definition of Model Fiber Orientations**

The model tissue had an elliptical profile, with major and minor radii that varied as a function of slice number. To create this profile, a mask signal, $S_M$, was calculated as:

$$S_M = {r^2}/{a(s)} + {c^2}/{b(s)} \qquad \text{[S1]}$$

With *r* the distance from the image center in pixels along the row dimension and *c* the distance from the image center in pixels along the column dimension. *a* and *b* were empirically set in a slice number-dependent manner:

$$a = \begin{cases} 12s & s \leq 25 \\ 300 - 15 * |25 - s| & s > 25 \end{cases} \qquad \text{[S2a]}$$

$$b = \begin{cases} 6s & s \leq 25 \\ 150 - 10 * |25 - s| & s > 25 \end{cases} \qquad \text{[S2b]}$$

with *s* the slice number. A composite tissue mask was formed by assuming values $S_M > 1$ to lie outside of the tissue boundaries and values $\leq 1$ to lie inside of the tissue boundaries. The aponeurosis was centrally located in the row and column directions, with a maximum size of three pixels in the column direction, a maximum size in the row direction that was eight pixels less than the corresponding muscle size for the given slice, and round anterior and posterior edges. The muscle boundary mask was defined as the subtraction of the aponeurosis boundary mask from the composite tissue mask.

To model the muscle fiber orientations, the in-plane (azimuthal) and through-plane (elevation) components were separately considered. The in-plane components were modeled as follows. Points defining the outer muscle boundary and the muscle aponeurosis boundary were identified using MATLAB's *bwboundaries( )* function. The minimum row/median column coordinates in each boundary were aligned. The vectors were resized to a length of 150 points and redundant points were eliminated. Corresponding points were connected using straight lines (Figure S1).

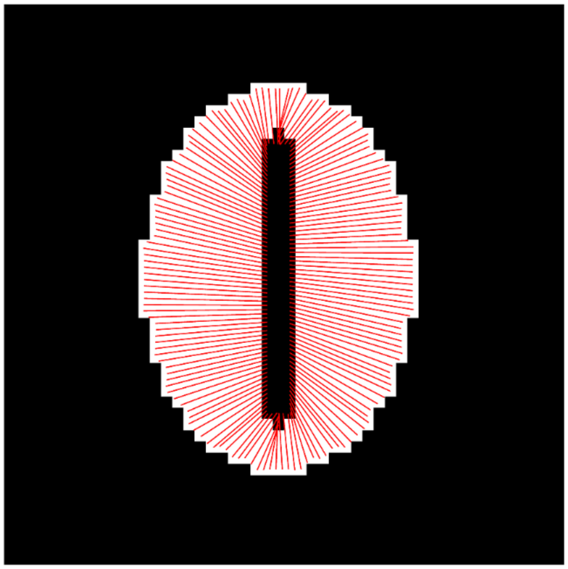

**Figure S1. Azimuthal muscle fiber orientations in the simulated muscle.** White portions of the image show pixels contained within the muscle boundary mask. Interior black pixels show the central aponeurosis. Red lines show in-plane muscle fiber orientations as lines connecting paired points along the aponeurosis and the outer muscle boundary.

The rise ($\Delta y$) and run ($\Delta x$) of each line was converted to a unit-length vector, [$\Delta x$ $\Delta y$], that was used to describe the in-plane fiber orientation. Expressed in a spherical coordinate system, $\Delta x$ and $\Delta y$ describe the azimuthal angle, $\phi$. To model the elevation angle $\theta$, the $\Delta z$ component was defined as:

$$\Delta z = cot\left(\theta_B - \left(L_S + {}^{L_S}/_{6}\right)^2 - {}^{s}/_{10}\right) \quad [S3]$$

With $\theta_B$ the base elevation angle (=25°) and $L_S$ the distance along the straight-line segment connecting the corresponding points. The second term on the right-hand side of Eq. S3 was set to cause $\theta$ to increase as a function of distance from the slice center, and the third term on the right-hand side of Eq. S3 was set to cause $\theta$ to increase as a function of slice number. This equation was solved at 20 linearly spaced points between the endpoints. At each intermediate point, the fiber orientation vector [$\Delta x$ $\Delta y$ $\Delta z$] was constructed, converted to unit length, and taken as the local first eigenvector of the diffusion tensor, $\varepsilon_1$.

The high resolution of points along the muscle boundaries, and the solution of Eq. S3 at 20 points along the path between the corresponding points, caused there to be voxels with overlapping model muscle fibers. To form a net $\varepsilon_1$ at each voxel, the $\varepsilon_{1x}$, $\varepsilon_{1y}$, and $\varepsilon_{1z}$ components of all muscle fibers passing through a voxel were separately summed, and the resulted summed vector was converted to unit length. Figure 1 in the main text illustrates the azimuthal and elevation angles corresponding to these net vectors.

### S2. Seed Point Illustrations

Figure S2 illustrates the points contributing to the restricted and expanded seeding meshes in the APO seeding methods.

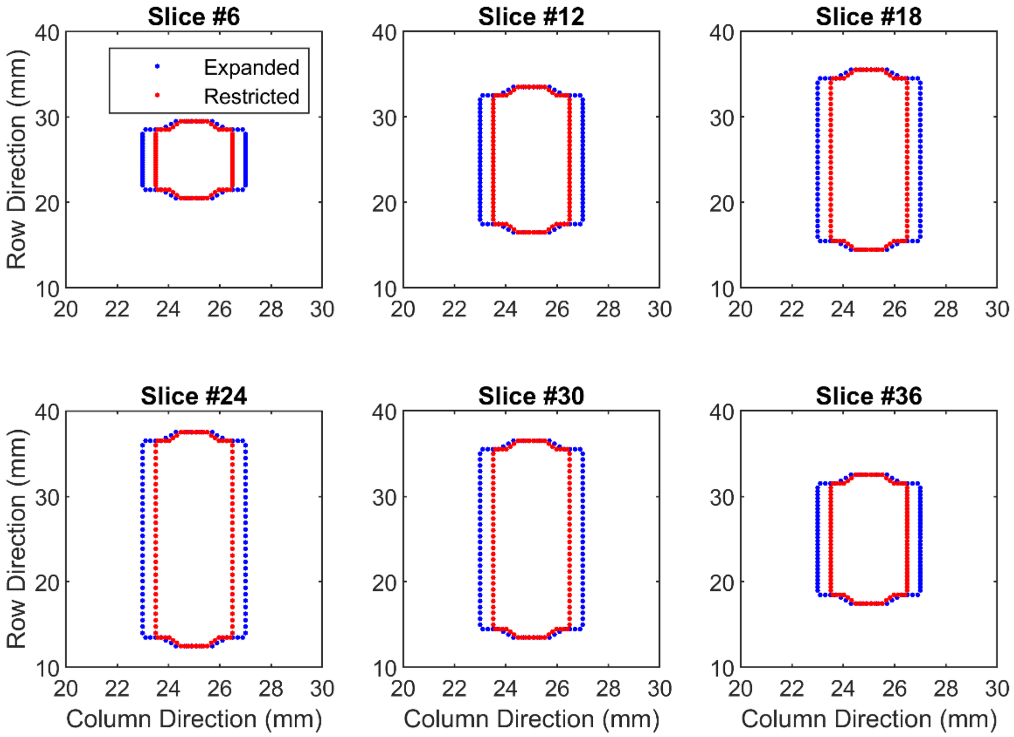


**Figure S2. Aponeurosis mesh points.** The restricted mesh fell 10 µm inside of the muscle-aponeurosis boundary, while the expanded mesh fell up to 0.5 mm outside of this boundary.

Figure S3 illustrates the seed point distributions for the VXL-1 condition.

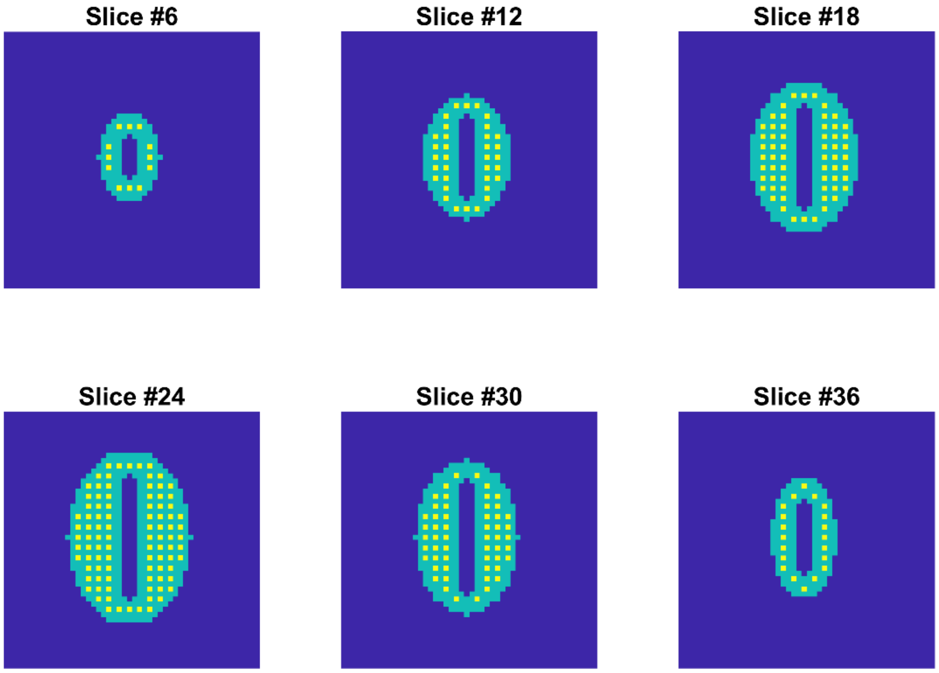


**Figure S3. Seed point distribution, VXL-1 condition.** The seed points were placed at a frequency of 1 point/2 voxels in the row and column directions, within a once-eroded muscle boundary mask. Dark blue portions of the images represent image regions outside of the muscle; yellow portions indicate seed points; and teal portions represent unseeded portions of the original muscle boundary mask.

Figure S4 illustrates the seed point distributions for the VXL-2 condition.

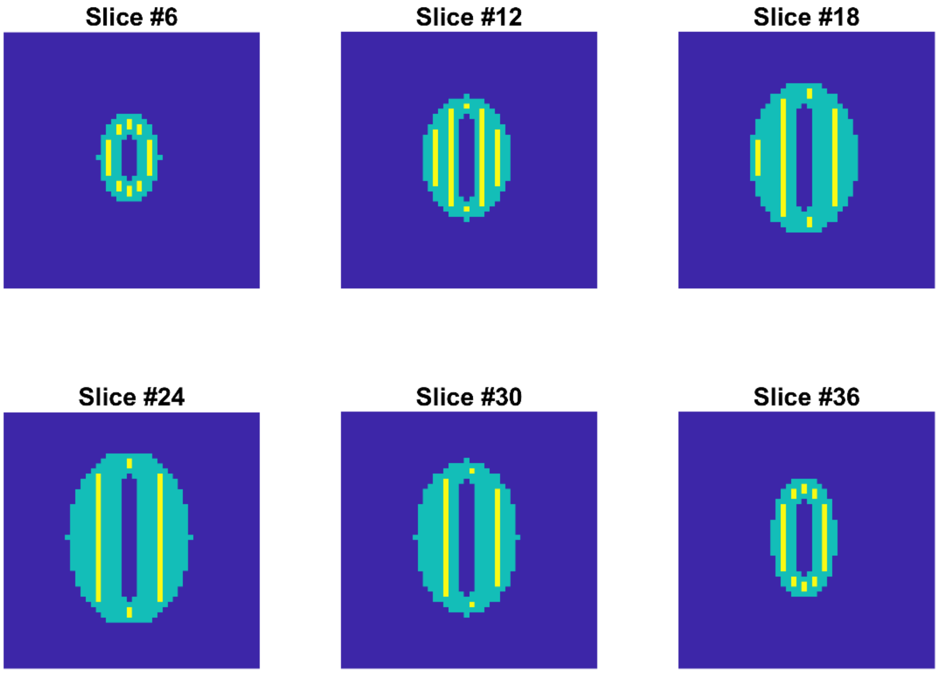


**Figure S4. Seed point distribution, VXL-2 condition.** The seed points were placed at a variable frequency in the column direction, depending on the size of the region of interest, within a once-eroded muscle boundary mask. Dark blue portions of the images represent image regions outside of the muscle; yellow portions indicate seed points; and teal portions represent unseeded portions of the original muscle boundary mask.

Figure S5 illustrates the seed point distributions for the VXL-3 condition.

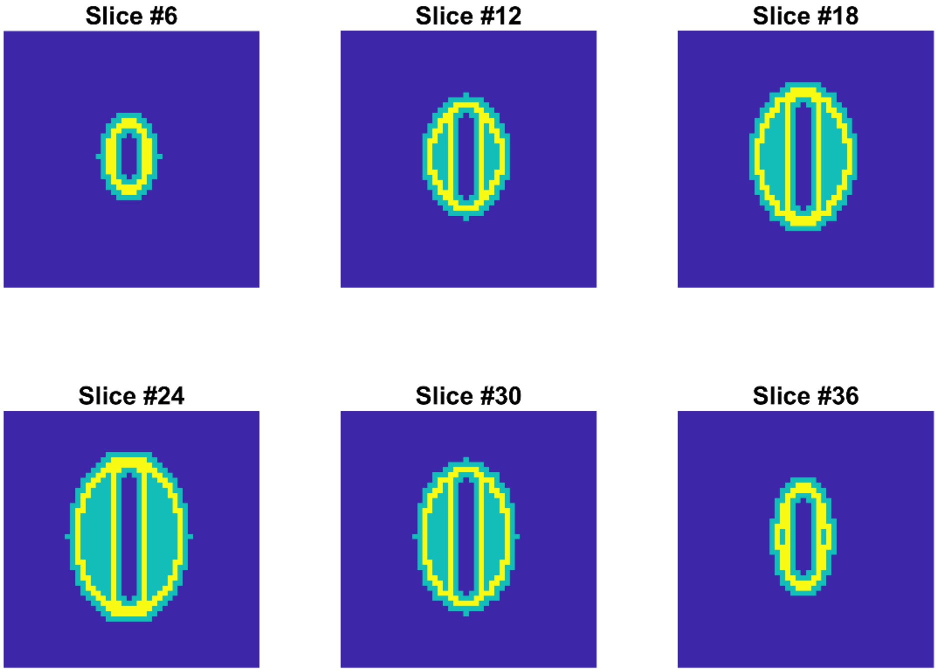


**Figure S5. Seed point distribution, VXL-3 condition.** The seed points were placed one voxel interior to the outer edge, and one voxel exterior to the central aponeurosis, of a once-eroded muscle boundary mask. Dark blue portions of the images represent image regions outside of the muscle; yellow portions indicate seed points; and teal portions represent unseeded portions of the original muscle boundary mask.

### S3. MuscleDTI_Toolbox Modifications

Table S1 provides a list of function versions employed and their modifications from previous publications.

**Table S1. MuscleDTI_Toolbox functions used in the present work.** Files are located at https://github.com/bdamon/MuscleDTI_Toolbox.

| Function | Purpose | Version | Modifications from Prior Versions |
|---|---|---|---|
| *define_roi( )* | Uses aponeurosis mask to define aponeurosis seeding mesh | 1.0.0 | No modifications |
| *fiber_track_v20( )* | Propagates fiber tracts from seed points | 2.0.0 | Introduces several options for voxel-based seeding; for voxel seeding methods, eliminates points lying in the internal aponeurosis; removes fiber assignment by continuous tractography as a tracking algorithm option |
| *fiber_smoother_v14( )* | Performs polynomial smoothing for originally propagated fiber tracts | 1.4.0 | Minor bug fixes; adds ability to weight initial point in tract more heavily, improving fit quality; ensures compatibility with new fiber tract matrices following creation of new seeding capabilities |
| *fiber_quantifier_v20( )* | Obtains estimates of fiber tract orientations, length, and curvature | 2.0.0 | Adds calculation of α, β, and γ angles; ensures compatibility with new fiber tract matrices following creation of new seeding capabilities |
| *fiber_visualizer_v11( )* | Visualizes images with aponeurosis mesh, muscle boundary mask, and/or fiber tracts | 1.1.0 | Provides correct units for axis length scales |
